# SIGLEC1 FACILITATES MACROPHAGE-CD8^+^ T CELL INTERACTIONS AND CORRELATES WITH CANCER IMMUNOTHERAPY RESPONSE

**DOI:** 10.64898/2026.07.09.737484

**Authors:** Sofía Ibáñez-Molero, Magali Coccimiglio, Babet Olivia Springer, Megan Bowien de Ruiter, Georgia Clayton, Steven Wijnen, Christian Blank, Mariette Labots, Tanja de Gruijl, Yvette van Kooyk

## Abstract

Antigen-presenting cell (APC) interactions with cytotoxic T cells are critical for anti-tumour immunity and response to immune checkpoint blockade (ICB), yet context-specific regulators in the tumour microenvironment remain not fully defined. Here we identify the lectin receptor SIGLEC1 as a key mediator of macrophage–T cell interactions in human melanoma. In a well-characterized patient cohort, SIGLEC1 was selectively upregulated in inflammatory macrophages physically associated with activated/exhausted CD8⁺ T cells. Imaging and functional analyses revealed that SIGLEC1 accumulates at the macrophage–T cell interface and promotes cell clustering. SIGLEC1 ligands were enriched on activated T cells, and their blockade reduced cytotoxic cytokine production *ex vivo*. Single-cell (spatial) transcriptomics across independent ICB-treated melanoma cohorts showed that SIGLEC1⁺ macrophages localize near CD8⁺ T cells and are enriched in responders, where they also associate with T cells expressing activation/exhaustion markers. These findings define a SIGLEC1-dependent macrophage–T cell niche linked to effective immunotherapy.

## INTRODUCTION

Immunotherapies, such as immune checkpoint blockade (ICB), have notably improved the clinical outcome of melanoma, yet, a subset of patients still does not show benefit^1^. Effective immunotherapy requires the infiltration of tumours by effector immune cells, including cytotoxic T cells and antigen presenting cells (APCs). Priming of T cells by APCs is essential for the acquisition of cytotoxic function, enabling their migration into the tumour microenvironment (TME) where they can recognize cognate cancer antigens and promote tumour cell killing. The abundance of APCs and T cells in tumours is a well-established predictive and prognostic markers of ICB response^2–7^. However, infiltration alone is insufficient; the phenotypic state, activation status and spatial distribution of immune cells within the TME are also critical determinants of therapy efficacy^8,9^.

APCs are a heterogeneous group of immune cells comprising macrophages, dendritic cells (DCs) and B cells, which process and present antigens to T cells, leading to their activation, often linked to effective ICB treatments. However, APCs can also act as suppressors of cytotoxic T cell activity. For example, CXCL9/10-producing inflammatory macrophages are associated with positive ICB responses^10^, whereas SPP1+ immunosuppressive macrophages correlate with poor clinical prognosis^11,12^. Thus, macrophage populations modulate antitumour immunity through direct stimulatory or inhibitory interactions with T cells. Prognostically, the abundance and activation state of infiltrating T cells is as important as APC infiltration^8^. Certain T cell exhausted or dysfunctional states have been linked to improved antitumour responses, whereas T cells in these states can also display reduced cytotoxic capacity^8,13,14^. T cell phenotypes and functionalities are shaped by interactions with neighboring cells, and thereby highly dependent on their spatial organization within the TME.

Recent advances in spatial transcriptomics have revealed that cell-cell proximity in tumours is a strong predictive marker for immunotherapy. For instance, inflammatory macrophages in close proximity to T cells correlate with positive ICB response^13^, whereas T cells near SPP1+ immunosuppressive macrophages are linked to unfavorable outcomes in multiple cancer types^7,12,15^. Furthermore, these immunosuppressive macrophages can sequester T cells at the tumour margin, restricting their access to cancer cells^16,17^. These observations underscore the importance of understanding not only the abundance of APC and T cells, but also how and where they communicate within the TME.

To better understand these interactions, we previously developed an approach to isolate and analyze physically interacting cell clusters from melanoma tumours^18^. Unlike earlier studies that inferred interactions from cell–cell proximity in two-dimensional histological sections, this method captures physically engaged cells from entire three-dimensional tumour biopsies. Single-cell RNA sequencing (scRNA-seq) revealed T cells in clusters with multiple APC subsets, including a population of inflammatory macrophages. Notably, when T cells derived from these macrophage–T-cell clusters were expanded using a rapid expansion protocol (REP), which disrupts macrophage contact, they displayed enhanced anti-tumour activity both *ex vivo* and *in vivo*. Here, we aimed to identify cell-surface receptors that distinguish macrophages that engage T cells from those that do not. This led to the identification of SIGLEC1, for which we subsequently assessed its functional role in macrophage-T-cell interactions, T cell activation as well as its spatial distribution in me lanoma tissues.

## RESULTS

### SIGLEC1 is higher expressed in inflammatory macrophages that are interacting with CD8+ T cells in tumours from melanoma patients

To investigate transcriptional differences between interacting and non-interacting APCs and CD8⁺T cells, we previously performed scRNA-seq of monocytes/macrophages (Mo/Mac) and CD8⁺ T cells isolated either as clusters or singlets from melanoma patients (n = 5) (**Fig. 1a**)^18^. Here, we performed differential expression analysis that revealed multiple genes upregulated in Mo/Macs in clusters with CD8⁺ T cells compared to non-interacting Mo/Macs (log2FC>0.5). Among these genes, we focused on cell surface receptors as they are potential mediators of these Mo/Macs-T cell interactions. We identified SIGLEC1 as the only upregulated protein in the interacting Mo/Macs that was also a cell surface receptor (**Fig. 1b**), which was particularly higher expressed in inflammatory macrophages (previously defined as C1q^hi^ inflammatory macrophages, defined by high expression of CXCL9-10^18^) (**Fig. 1c**).

**Figure 1.**
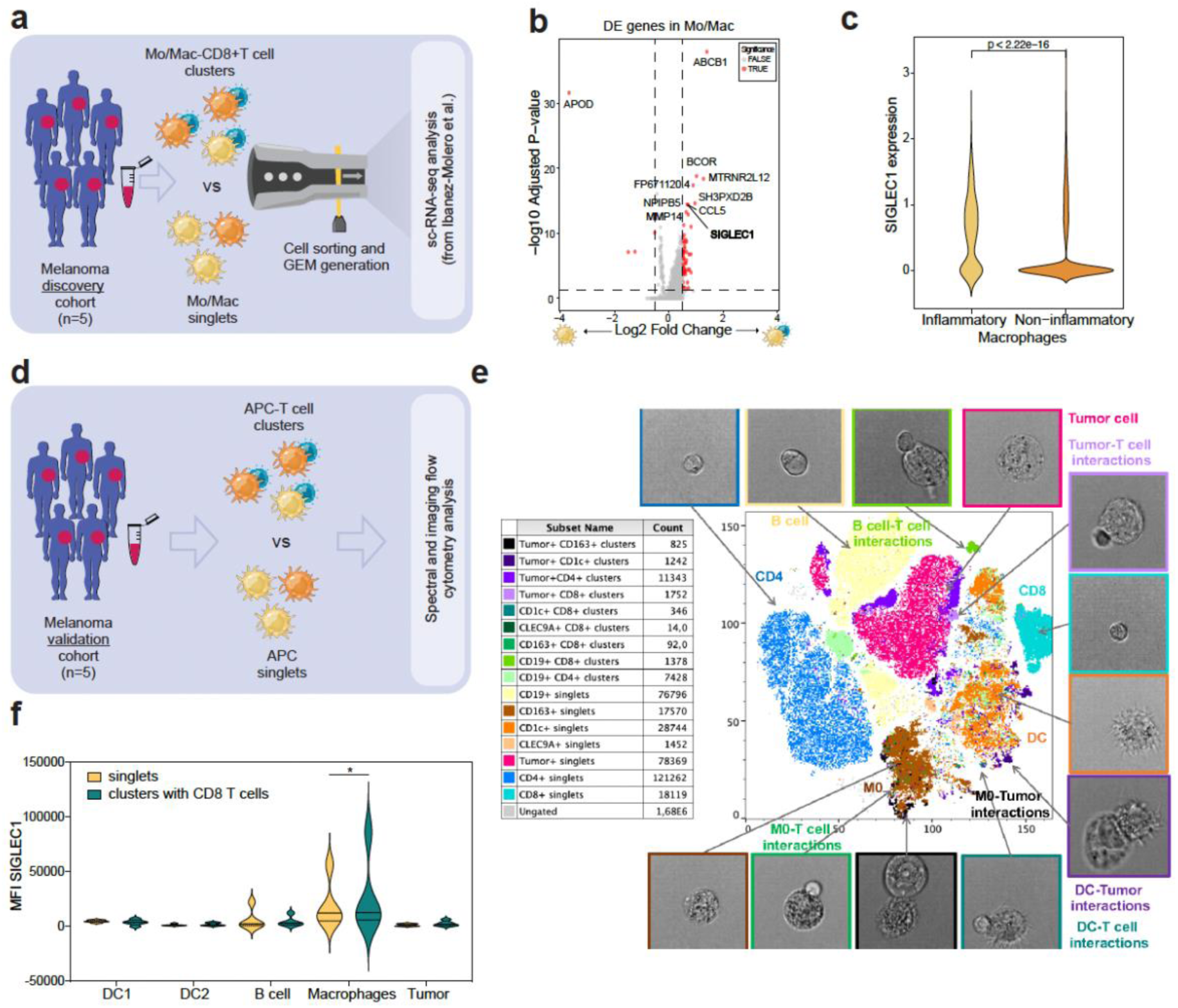
Single-cell differential gene expression analysis reveals higher expression of SIGLEC1 in inflammatory macrophages that are in cluster with CD8+ T cells. a) Schematic representation of discovery patient cohort analysis. b) Differential gene expression analysis in Mo/Mac clusters versus singlets. Significant genes are indicated as red dots. Top genes are outlined. c) SIGLEC1 mRNA expression in Mo/Mac inflammatory and non-inflammatory defined in Ibáñez-Molero, Veldman et al.^18^ Wilcox test used for statistical analysis d) Schematic representation of validation patient cohort analysis. e) tSNE analysis and representative pictures of singlets and clusters found in a representative melanoma patient. Four additional patients shown in Extended Data Fig. 1a. f) SIGLEC1 expression in different APC subtypes as singlets or clusters with CD8+ T cells. 2way ANOVA test used for statistical analysis.

To validate these mRNA-based findings at the protein level and to assess SIGLEC1 expression across additional APC subsets, we analyzed a second cohort of melanoma patients (n = 5) using imaging flow cytometry and a newly developed spectral flow cytometry panel incorporating T cell and APC markers (**Fig. 1d**). This approach enabled identification of multiple APC populations, including DCs, B cells, and macrophages, present either as singlets or in physical association with T cells (**Fig. 1e, Supp Fig. 1a**). SIGLEC1 protein expression was significantly elevated in macrophages clustered with CD8⁺ T cells—whereas no significant differences were observed in other APC subsets (**Fig. 1f, Extended Data Fig. 1b**). In parallel, as control, we evaluated two additional SIGLEC family members, SIGLEC7 and SIGLEC9, neither of which showed comparable enrichment in macrophages (**Extended Data Fig. 1c**). Together, these data demonstrate that SIGLEC1 is upregulated at both the mRNA and protein levels in macrophages physically interacting with CD8⁺ T cells in human melanoma tumours.

### SIGLEC1 re-localizes to the immune interface between inflammatory macrophages and CD8+ T cells and facilitates their cluster formation

SIGLEC1 (also known as CD169) is an immunoglobulin-like lectin receptor expressed on inflammatory macrophages that binds sialylated cell surface glycans^19,20^. Upon antigen recognition, APCs and T cells form an immunological synapse, defined by re-localization of membrane receptors and reduced T cell motility^21–25^. Given the enrichment of SIGLEC1 in macrophages engaged with CD8⁺ T cells, we examined whether it similarly re-localized to the synapse to stabilize these interactions.

Monocytes isolated from healthy donor peripheral blood mononuclear cells (PBMCs) were polarized into inflammatory and suppressive macrophages, or non-polarized; and subsequently co-cultured with autologous CD8⁺ T cells for 1 hour (**Fig. 2a**). Localization of SIGLEC1 was assessed by imaging flow cytometry (**Fig. 2b, Extended Data Fig. 2a, b**). Consistent with our patient-derived data, inflammatory macrophages exhibited higher baseline SIGLEC1 expression (**Fig. 2c**). Upon co-culture with CD8⁺ T cells, SIGLEC1 in inflammatory macrophages re-localized to the intercellular interface, whereas no such re-localization was observed in suppressor or non-polarized macrophages (**Fig. 2d**). These findings indicate that SIGLEC1 is recruited to the immune synapse between inflammatory macrophages and CD8⁺ T cells, suggesting a potential role in synapse formation and/or maintenance.

**Figure 2.**
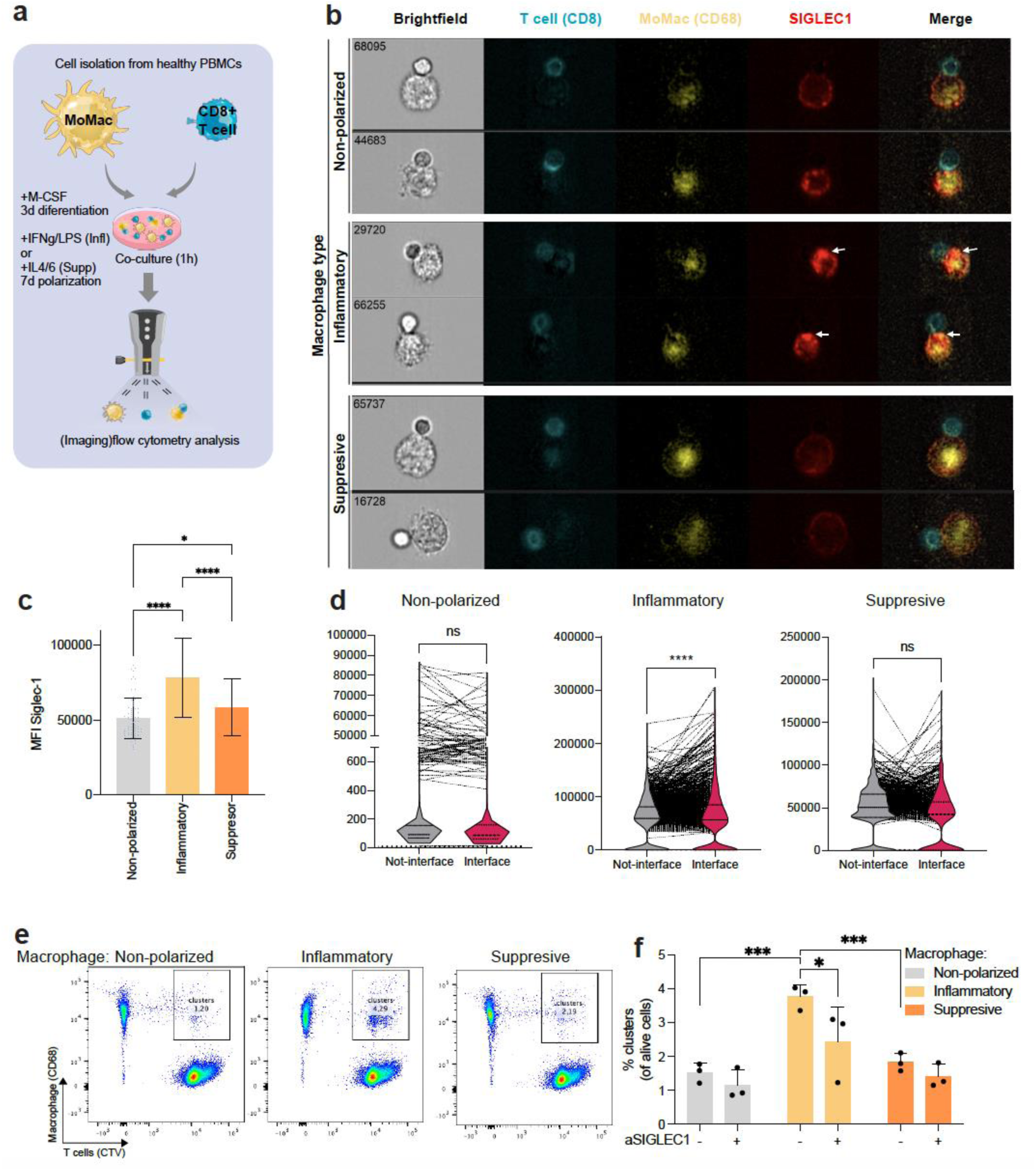
SIGLEC1 re-localizes to the immune interface between inflammatory macrophages and CD8+ T cells and facilitates their cluster formation. a) Schematic representation of monocyte-derived macrophage-T cell *in vitro* co-culture model. b) Imagestream analysis of co-cultures from (a) after 1h incubation. Three types of macrophages are used: non-polarized, inflammatory and suppressive. Markers used are indicated in colors: blue (CD8), yellow (CD68) and red (SIGLEC1). Arrows indicate interface between two cells with SIGLEC1 re-localization. c) Median fluorescence intensity (MFI) of SIGLEC1 in macrophages from (b). Ordinary one-way ANOVA used for statistical test. d) Interface re-localization analysis performed on the three macrophages subtypes from (b). Lines indicate individual cells expression at their interface or not-interface. Paired t-test used for statistical analysis. e) Representative flow cytometry plots of macrophage-T cell co-cultures, gated on double positive events as indicated (i.e. clusters). Number on the gate indicate the frequency of clusters from alive cells f) Cluster formation quantification from (e) with anti-SIGLEC1 (+) or isotype (-) treatments. 2way ANOVA test used for statistical analysis.

Next, we assessed whether SIGLEC1 contributed to maintaining macrophage-T cell interaction. Macrophage–T cell co-cultures were treated with SIGLEC1-blocking antibody, and cluster formation was quantified as a measure of cell–cell interaction. Inflammatory macrophages formed the highest proportion of clusters, confirming our patient-derived observations (**Fig. 2e, f**). Upon SIGLEC1 blocking, macrophage-T cell clusters were reduced for inflammatory macrophages, but no significant differences were detected in suppressor or non-polarized macrophages, likely due to their low baseline interaction frequencies (**Fig. 2e, f**). These results indicate that SIGLEC1 promotes stable contacts between inflammatory macrophages and CD8⁺ T cells.

### SIGLEC1 ligands are higher expressed in activated T cells and their blockade *ex vivo* reduces T cell cytokine production

In our previous work we showed that CD8⁺ T cells within Mo/Mac–T cell clusters predominantly exhibited activated or exhausted transcriptional states (**Fig. 3a**). Thus, we hypothesized that activated CD8⁺T cells should be more capable of engaging Mo/Mac via increased expression of SIGLEC1 ligands. We examined the expression of SIGLEC1 ligands on CD8⁺T cells *ex vivo* in single-cell–dissociated melanoma tumours and its association with CD8⁺T cell activation, as evaluated by CD137 and PD-1 expression. We showed that CD8⁺ T cells expressing SIGLEC1 ligands displayed higher levels of activation markers (**Fig. 3b**). Because macrophages also express SIGLEC1 ligands, we could not selectively assess ligand expression on CD8⁺T cells within clusters versus singlets. Nonetheless, these data indicate that SIGLEC1 ligand expression correlates with higher T cell activation.

**Figure 3.**
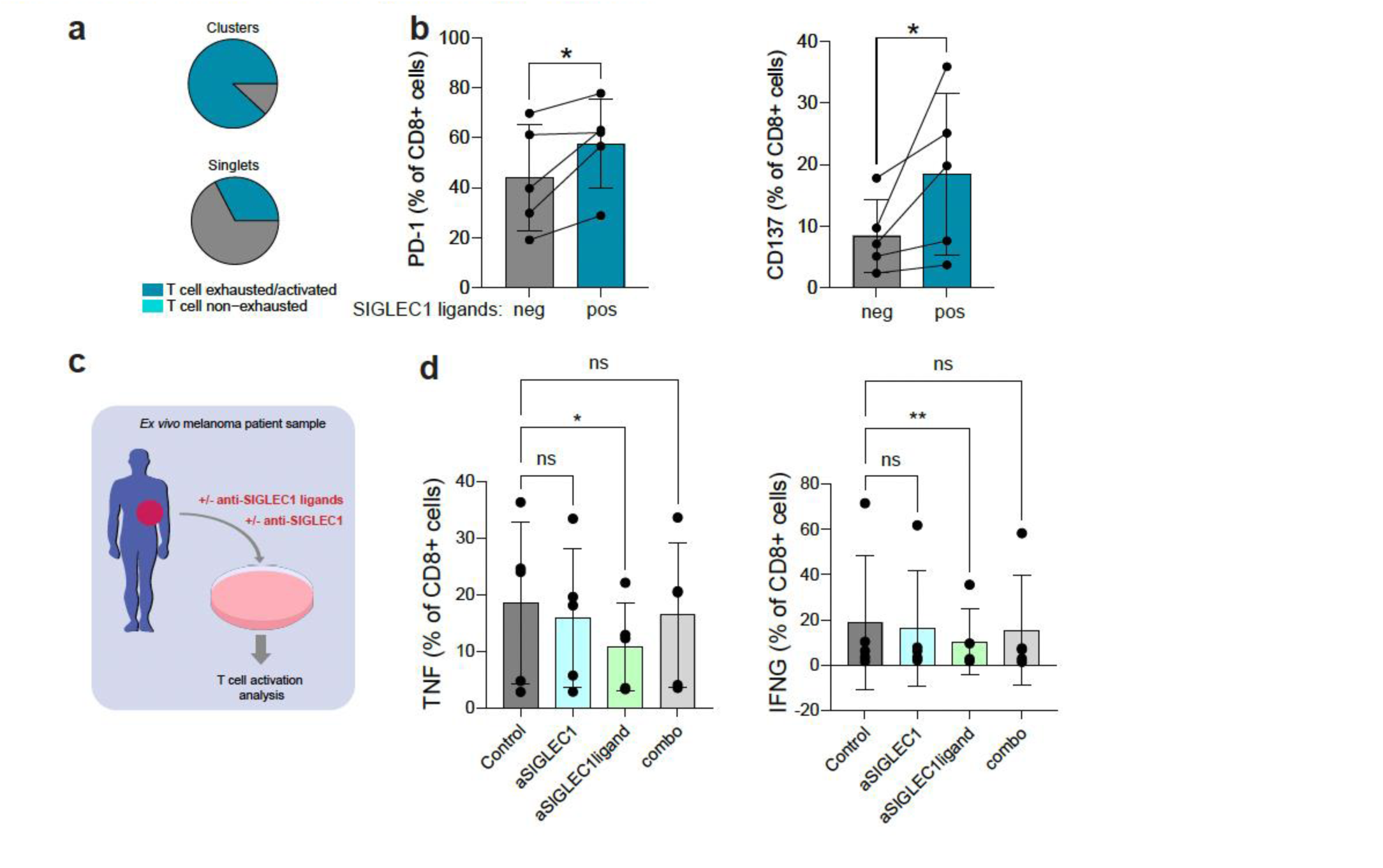
SIGLEC1 ligands are higher expressed in activated T cells and their blockade *ex vivo* reduces their cytokine production. a) Frequencies of exhausted/activated and non-exhausted CD8^+^T cells singlets and clusters from scRNA-seq cohort from Ibáñez-Molero, Veldman et al. b) Frequency of T cells positive for activation markers (PD1 and CD137) in populations negative (neg) and positive (pos) for SIGLEC1 ligand expression, measured with mouse recombinant SIGLEC1-fc construct. Ratio paired t test used for statistical analysis. c) Schematic representation of *ex vivo* melanoma patient sample experiment, treated with anti-SIGLEC1 blocking antibody or anti-SIGLEC1 ligands blocking, or combination (combo) for four days. d) Quantification of frequency of TNF and IFNG positive CD8^+^ T cells from (c). Lognormal RM one-way ANOVA test used for statistical analysis.

To assess the functional role of SIGLEC1 in CD8⁺ T cell activation, we treated *ex-vivo* single-cell–dissociated melanoma tumours with a blocking antibody against SIGLEC1, and SIGLEC1 recombinant protein that blocks SIGLEC1 ligands, or both in combination, which is expected to neutralize their reciprocal interaction (**Fig. 3c, Extended Data Fig. 3c, d**). Blockade of SIGLEC1 ligands resulted in reduced TNF and IFNγ production by CD8⁺ T cells, whereas SIGLEC1 blockade alone had no significant effect, suggesting additional functions of SIGLEC1-ligands axes within the TME (**Fig. 3d**). No significant changes were observed in T cell surface activation markers (**Extended Data Fig. 3a**). Together, these results indicate that SIGLEC1 ligand expression correlates with T cell activation and that ligand blockade modulates CD8⁺ T cell cytotoxic cytokine production.

### SIGLEC1 expression in macrophages and their proximity to T cells predict ICB response

Spatial context is critical, as the proximity of immune subsets influences possible cell–cell communication and therapeutic response, therefore we examined SIGLEC1 expression and the spatial distribution of macrophages and T cells *in situ*. We performed scRNA-seq and spatial transcriptomics on two independent melanoma cohorts treated with ICB, for which we analyzed paired pre- and post-treatment biopsies when available (**Fig. 4a**, Coccimiglio, Ibáñez-Molero et al. under revision). SIGLEC1 expression was primarily restricted to macrophages and DCs, consistent with its established role in myeloid populations (**Fig. 4b**). In the first cohort, differential expression analysis on macrophages revealed SIGLEC1 as one of the most significantly enriched genes in macrophages from responders compared to non-responders (**Fig. 4c**). We could validate this finding also in our second cohort (**Fig. 4d**).

**Figure 4.**
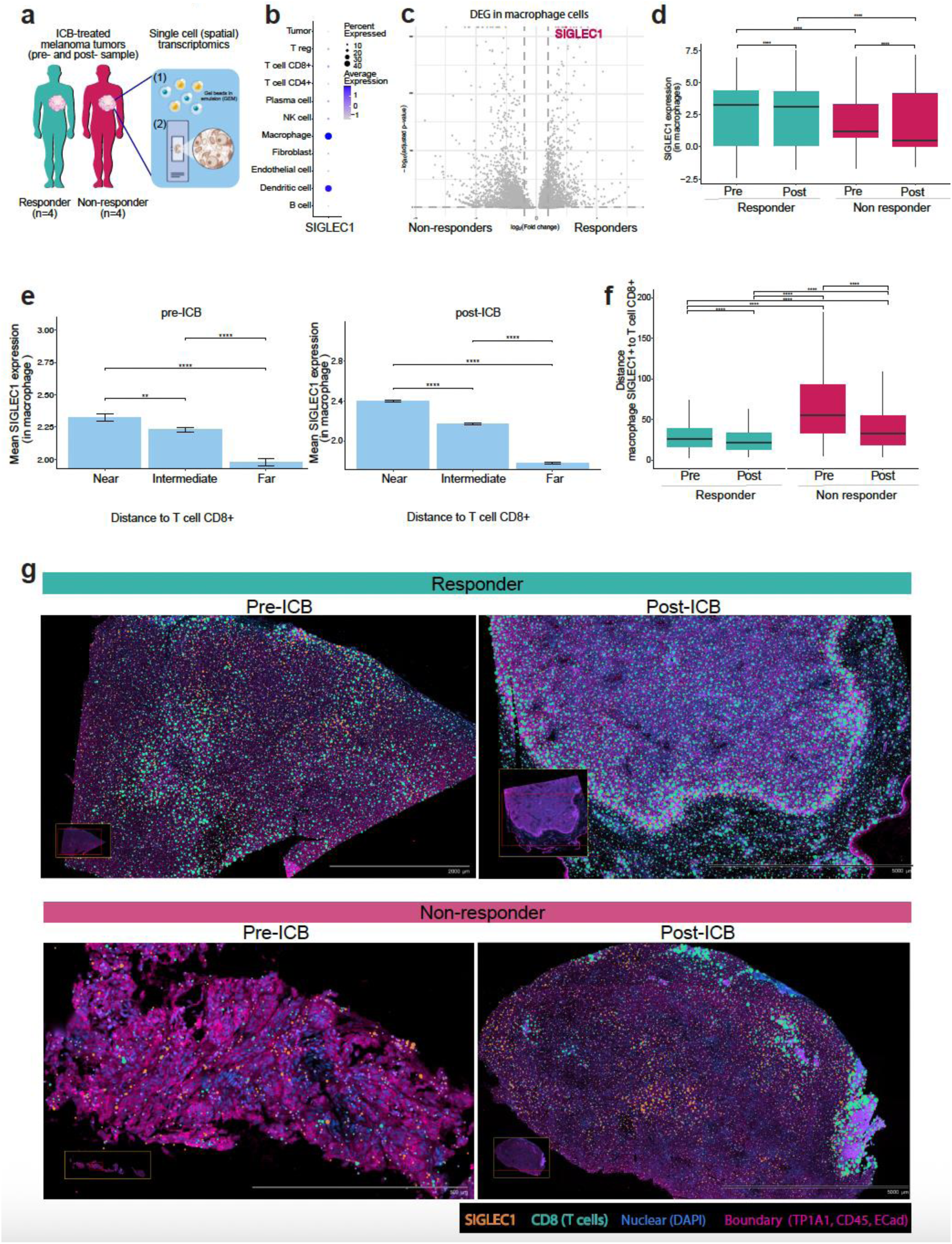
SIGLEC1 expression on macrophages and their proximity to CD8+ T cells predict ICB response. a) Schematic representation of spatial transcriptomics ICB-treated melanoma patient cohorts (n=4, n=4). In the first cohort four pre-ICB melanoma samples were included (two responders and two non-responders) and analyzed by scRNA-seq. In the second cohort three matched ICB-treated melanoma patient samples pre- and post-ICB, and one patient only pre- ICB were included and analyzed for spatial transcriptomics (Xenium), a total of seven tumour tissue slides, as described in Coccimiglio, Ibáñez-Molero et al. (under revision). b) SIGLEC1 mRNA expression in all cell types annotated in the second cohort. c) Differential gene expression analysis between responders and non-responders, performed on macrophage cells from the first cohort. Dotted lines indicate P<0.05 and log2 (fold change)>0.5. d) SIGLEC1 mRNA expression in macrophage cells from second cohort in responders and non-responders, pre-and post-ICB treatment on the second cohort. Wilcox with Bonferroni correction test used for statistical analysis. e) Mean SIGLEC1 expression in macrophage cells relative to distance groups from the closest target cell. “Near” corresponds to the first tercile, “Intermediate” to the second tercile, and “Far” to the third tercile of all distances calculated. Bar plots show the average SIGLEC1 expression per group with standard deviation. Wilcoxon test with Bonferroni correction used for statistical analysis. Analysis performed on pre-ICB samples (left) and post-ICB samples (right). f) Distance of SIGLEC1+ macrophage cells to CD8+ T cells in responders and non-responders pre- and post-ICB treatment. Wilcox test used for statistical analysis. g) Example pictures of responder and non-responders, pre- and post-ICB patient samples Dots depict expression of genes: SIGLEC1 (orange), CD8 (blue, T cell marker).

Leveraging the spatially resolved dataset, we next assessed SIGLEC1 expression relative to macrophage proximity to CD8⁺ T cells. Macrophages located near CD8⁺ T cells displayed markedly higher SIGLEC1 levels than those positioned farther away, both before and after ICB treatment (**Fig. 4e**). Additionally, given that SIGLEC1 was also expressed in DCs, we evaluated the correlation of SIGLEC1 expression on DCs with their distance to T cells, showing an opposite trend to macrophages (**Extended Data Fig. 3b**). The distance between SIGLEC1⁺ macrophages and CD8⁺ T cells was smaller in responders compared to non-responders. Upon ICB treatment, this distance was decreased in both, responders and non-responders (**Fig. 4f, g**). Together, these findings indicate that elevated SIGLEC1 expression in macrophages and their close spatial association with CD8⁺ T cells define immune niches associated with favorable ICB outcome.

### SIGLEC1 expression on macrophages correlates with T cell activation and exhaustion signatures that differentiate ICB outcomes

We initially identified SIGLEC1 as a gene enriched in macrophages interacting with exhausted or activated T cells (**Fig. 1**, **Fig. 3a** and described in Ibáñez-Molero, Veldman et al.). Therefore, we investigated whether the presence of exhausted/activated T cells correlated with SIGLEC1⁺ macrophages in the TME in our melanoma cohort. We assessed the expression of canonical activation and exhaustion markers (LAG3, PRF1, GZMH, GZMA, GZMB, TOX, GZMK, TNF and PDCD1) alongside SIGLEC1 levels in macrophages. SIGLEC1 expression in macrophages strongly correlated with T cell activation and exhaustion signatures, specifically with TOX, GZMK, TNF and PDCD1 expression on CD8⁺ T cells (**Fig. 5a, b**). Both SIGLEC1 and T cell activation markers were elevated in responders compared to non-responders before treatment and most were further increased after ICB treatment (**Fig. 5b**). Although we focused our analysis on CD8⁺ T cells, the trends observed were also extended to other cytotoxic cells such as NK and CD4⁺ T cells (**Extended Data Fig. 3c**). Additionally, SIGLEC1 expression on DCs, also correlated with ICB response, suggesting a broader role (**Extended Data Fig. 3c**). Collectively, these data link SIGLEC1 expression on macrophages with the activation state of CD8⁺ T cells in the TME and highlight its potential as a determinant of effective antitumour immunity and response to ICB.

**Figure 5.**
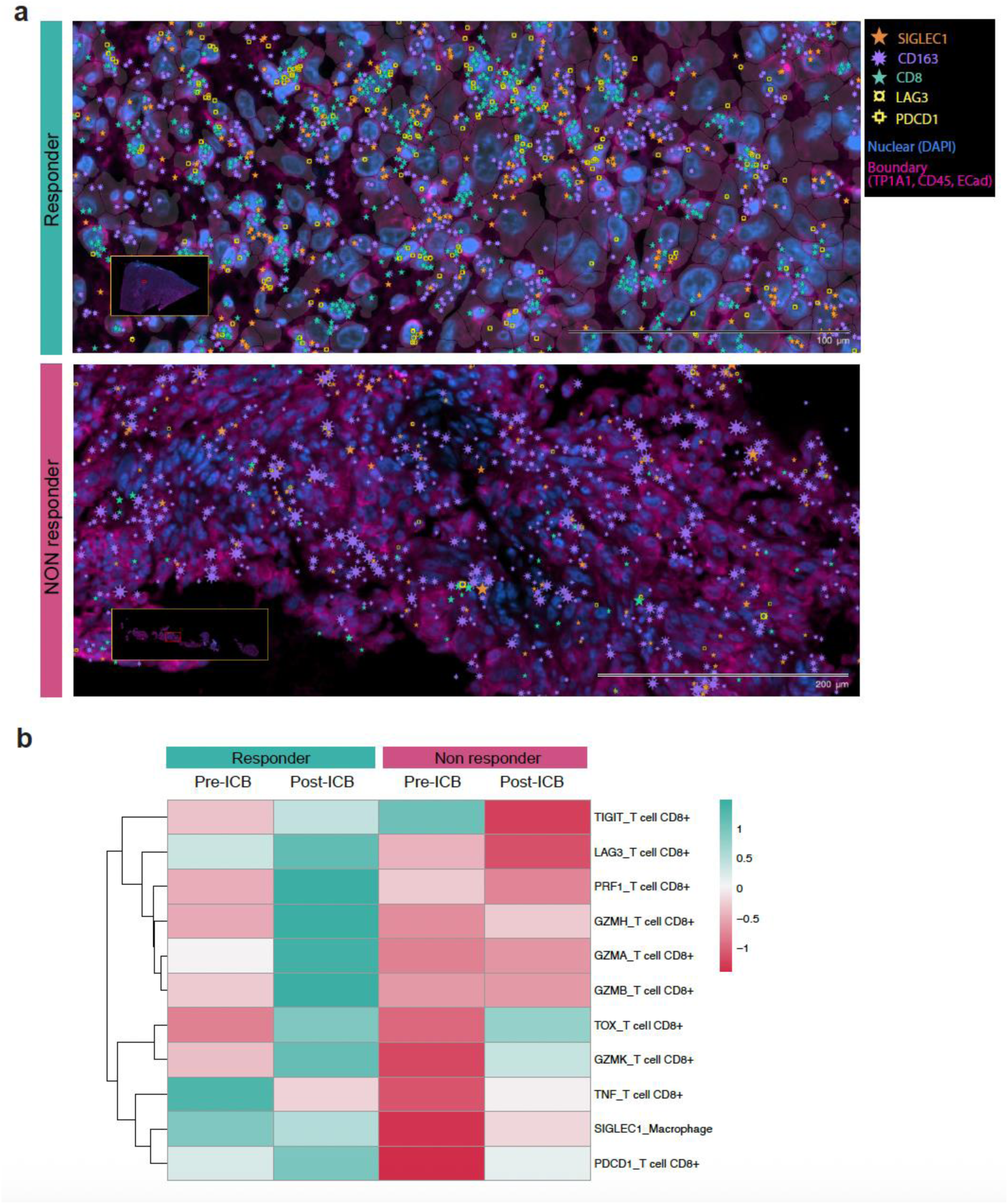
SIGLEC1 expression on macrophages correlates with T cell activation and exhaustion signatures that differentiate ICB response. a) Example pictures of responder and non-responders. Icons depict expression of genes: SIGLEC1 (orange), CD8 (blue, T cell marker), CD163 (purple, macrophage marker), LAG3 (yellow, ICB), PDCD1 (yellow, ICB). b) Heatmap showing gene expression of the indicated gene in the indicated cell type (eg. SIGLEC1 gene expression in macrophage cells). Unsupervised row clustered. Blue indicates higher relative expression, rede indicates relative lower expression.

## DISCUSSION

Effective ICB response depends not only on immune cell infiltration into tumours but also on their functional state, spatial organization, and quality of interactions between immune populations within the TME. By isolating physically engaged immune cell clusters from intact tumour biopsies, we foudn that SIGLEC1 is selectively upregulated in inflammatory macrophages interacting with CD8⁺ T cells. Mechanistically, we show that SIGLEC1 redistributes to the immune interface between inflammatory macrophages and CD8⁺ T cells, consistent with features of an immunological synapse. Blockade of SIGLEC1 reduced macrophage–T cell cluster formation, supporting a role for SIGLEC1 in stabilizing cell–cell contact. In patients, SIGLEC1 ligands are preferentially expressed on activated/exhausted T cells, and the spatial proximity of SIGLEC1⁺ macrophages to T cells predicts clinical response to ICB.

SIGLEC1 has been primarily studied for its role in antigen capture and presentation, particularly in lymphoid tissues, and has been leveraged for antigen-targeting vaccine strategies^26,27^. However, its function in mediating direct APC–T cell interactions within tumours has remained poorly defined. Given its extended extracellular domain and affinity for sialylated glycans, SIGLEC1 is well suited to mediate glycan-dependent adhesion, potentially prolonging cell interactions^28,29^.

We found that SIGLEC1 ligand expression was associated with T cell activation/exhaustion, defined by cell surface markers including CD137 and PD-1. These data suggest that sialylation patterns on activated T cells may promote engagement with SIGLEC1⁺ macrophages, reinforcing physical interactions within immune niches. Such interactions may facilitate sustained antigen presentation but could also contribute to chronic stimulation and the emergence of exhausted T cell states. This is in line with recent findings that show that the sialylation profile of T cells can determine their phenotype and functions^30^.

Functional perturbation experiments further highlight the complexity of the SIGLEC1–ligand axis. Blockade of SIGLEC1 ligands, but not SIGLEC1 itself, reduced TNF and IFNγ production by CD8⁺ T cells *ex vivo*. One interpretation is that SIGLEC1 ligands exert functions beyond binding SIGLEC1, potentially engaging additional glycan-binding receptors and influencing anti-tumour T cell functions^31^. Alternatively, SIGLEC1 blockade may interfere with other functions such as antigen uptake by macrophages, indirectly dampening T cell effector function^32,33^. Of note, we experimentally blocked a specific glycosylated motif by the use of a recombinant SIGLEC1 protein, but it is still not fully described which molecules carry this motif. The specific identity of the SIGLEC1 ligands remains unknown, and defining these ligands will be critical for understanding how glycan-mediated interactions regulate T cell responses in tumours.

Our single-cell (spatial) transcriptomic analyzes across two independent melanoma cohorts underscore the clinical relevance of SIGLEC1. SIGLEC1 expression in macrophages was enriched in patients responding to ICB, consistent with prior reports linking SIGLEC1⁺ macrophages to favorable outcomes in other malignancies, including endometrial and colorectal carcinomas^34,35^. Spatial analyzes further revealed that SIGLEC1-expressing macrophages preferentially localized in close proximity to CD8⁺ T cells and were correlated with T cells with activation and exhaustion gene signatures. Together, these findings support a model in which SIGLEC1–ligand interactions delineate immune niches characterized by sustained antigen engagement and active anti-tumour immunity.

Although our analyzes focused on macrophages, SIGLEC1 expression on other APC subsets, including DCs, also correlated with cytotoxic immune activation and ICB response, indicating that SIGLEC1–ligand interactions may broadly influence APC–lymphocyte communication^36^. However, SIGLEC1 expression on DCs anti-correlated with their distance to CD8^+^ T cells, an opposite finding than macrophages. Additionally, SIGLEC1 ligands were detected on macrophages themselves, raising the possibility of autologous or APC–APC interactions that could regulate antigen uptake, retention, or spatial organization within the TME. Dissecting these autologous roles will be important for understanding the full spectrum of SIGLEC1 function in tumours.

In summary, our study identifies the SIGLEC1–ligand axis as a regulator of macrophage–T cell interactions that coordinates spatial immune organization, T cell activation and exhaustion, and clinical response to ICB. By integrating single-cell, spatial, and functional analyzes, we reveal how glycan-mediated interactions contribute to the formation of immune niches that support antitumour immunity while we may still need checkpoint-mediated reinvigoration. These findings highlight SIGLEC1 and its ligands as potential biomarkers and modulators of immunotherapy responsiveness in cancer, which merit further investigation.

## MATERIAL AND METHODS

### Patient samples

Patients were enrolled under written informed consent in an institutional review board-approved clinical study of autologous whole-cell vaccination at the Amsterdam UMC, location VUmc (Amsterdam, The Netherlands)^37^. Resected tumour material (metastatic melanoma) was collected from melanoma patients undergoing surgery. All patients provided prior informed consent to research usage of material not required for diagnostics.

### T cells and monocyte isolation from PBMCs

PBMCs were isolated by density gradient with Ficoll-Paque PLUS (GE Healthcare, GE17-1440-02) from healthy donor buffy coats. CD14+ monocytes were isolated using CD14 Microbeads (Miltenyi, CAT 130-097-052) and T cells were isolated using CD8+ untouched beads isolation kit (Miltenyi, CAT 130-096-495). Monocytes were cultured in complete medium: RPMI1640 (Gibco) containing 10% Fetal Calf Serum (FCS, Biowest), 2mM L-Glutamine (Gibco) and 1000U per ml Penicillin-Streptomycin (Gibco). T cells were frozen after isolation. After thawing, T cells were cultured in RPMI1640 (Gibco) containing 10% Fetal Bovine Serum (FBS, Biowest), 2mM L-Glutamine (Gibco) and 1000U per ml Penicillin-Streptomycin (Gibco) and IL2 (10IU).

### Monocyte-derived macrophages differentiation and polarization

After isolation, CD14+ monocytes were differentiated into monocyte-derived macrophages (moMacs) for 6 days adding 50ng/ml M-CSF (Miltenyi, CAT 130-096-492) to complete monocyte medium (see above). Four days after isolation, moMacs are polarized into inflammatory or suppressor moMacs. For inflammatory macrophages IFNy (25ng/ml, Immunotools, CAT 11343536) and LPS (10ng/ml, Invivogen, tlrl-eblps) were added to the differentiation medium. For suppressor macrophages IL-4 (25ng/ml, Immunotools, CAT 11340047) and IL-6 (25ng/ml, R&D systems, CAT 206-IL-010) were added to the differentiation medium. Experiments are performed after two or three days of polarization.

### In vitro T cell: macrophage co-cultures

MoMac were harvested incubating them with TrypLE (Thermofisher, CAT 12604021) for 20min and T cells were thawed. Cells were added at a 1:1 ratio (100k T cells :100K moMacs) in non-adherent 96-well plates and incubated for 1h. For SIGLEC-1 blocking experiments, T cells were pre-stained with CTV (Invitrogen, CAT: C34557) for 20min at 37C. MoMacs were pre-treated with anti-SIGLEC1 (produced in house) or isotype antibodies (Ultra-LEAF™ Purified Mouse IgG1, κ Isotype Ctrl Antibody, Biolegend, CAT 400166) at 10ug/ml for at least 30min. After co-culture incubation, cells were resuspended and prepared for (imaging)-flow cytometry analysis.

### Flow cytometry analysis

Cells were washed with 0.1% bovine serum albumin (BSA) in phosphate-buffered saline (PBS). For surface staining, cells were stained with the indicated antibodies diluted in 0.1% BSA in PBS for 30 min on ice in the dark. After staining, cells were washed twice with 0.1% BSA in PBS and measured on a BD Fortessa flow cytometer. Data was analyzed using FACSDiva during acquisition and Flowjo for post-analysis.

For analysis of the patient validation cohort, cells suspensions were thawed and stained following the standard procedure above. Samples were then measured on an Aurora spectral flow cytometer.

### Imaging-flow cytometry analysis

Cells were stained following a standard flow cytometry protocol described in the previous section. In the final step, cells were resuspended at 10×10^6^ cells/ml in 0.1% BSA in PBS for measurement in CytPix or Imagestream® flow cytometers. For CytPix experiments (**Fig. 1**) cells were analyzed using Attune Cytometry Software. For Imagestream experiments (**Fig. 2**) data was analyzed using ImageStream® Mark II with INSPIRE acquisitions software during measurement and IDEAS® software for post-analysis. Immune synapse re-localization analysis was performed as described previously^18^.

### Patient samples ex vivo experiments

Tumor material was digested and frozen after resection. After thawing, melanoma cells were cultured at 500K in a 48 flat bottom wells-plate (Greiner Bio-One, CAT 677-180) in complete RPMI160 medium (see above) supplemented with IL2 (10IU). For a baseline readout 0.75 million melanoma cells were stained with flow panel 1 (described below). Simultaneously, melanoma cells were treated for four days at 37C with anti-SIGLEC1, isotype antibody (10µG/ML), Siglec1fc (1.8 µg/ml, produced in house) or a combination of anti-SIGLEC1 and Siglec1fc. After four days of incubation, cells were incubated with Brefeldin (InvivoGen, CAT inh-bfa) for 3 hours at 4C. Subsequently, melanoma cells were harvested by incubating them with TrypLE for 20min at 37C and resuspended and divided in two 96 v-bottom well plates (Sarstedt, CAT 82-1583) for flow cytometry. All cells were washed with 0.1% BSA in PBS. For intracellular staining, melanoma cells were stained with the indicated antibodies diluted in 0.1% BSA in PBS for 30 min on ice in the dark. The following antibodies were used for flow panel 1: LDnearIR-FVD780, FC-block, True monocyte stain, CD163-BV785, CD8-PE, NGFR/CD146-APC, CD137-PECy7, PD1-BV605 and Siglec-1-BV421 (see **Table 1**). The following antibodies were used for flow panel 2: LDnearIR-FVD780, FC-block, True monocyte stain, CD8-BV421, CD163-BV785, and Siglec-1-AF647 (produced in house) (see table). After staining cells were washed once with 0.1% BSA in PBS and once with 0.5% BSA in Hanks Balanced Salt Solution (HBSS). Subsequently, melanoma cells of both panels were stained with Siglec-1-FC and Siglec-1-Fc-AF488 (see **Table 1**) in 0.5% BSA in HBSS for 30 min on ice in the dark. After staining, cells were washed once with 0.5% BSA in HBSS and once with 0.1% BSA in PBS. Cells from flow panel 1 were measured in BD Fortessa flow cytometer. After washing, cells from panel 2 were fixed with 2% Paraformaldehyde (PFA) for 15 min on ice in the dark, followed by two washes of 0.5% saponin in 0.5% BSA in PBS. For intracellular staining, the cells were stained with the indicated antibodies dilute in 0.5% saponin in 0.5% BSA in PBS for 30 min on ice in the dark. The following antibodies were used: IFNg-PE and TNF-PerCP (see **Table 1**). After staining, the cells were washed once with 0.5% saponin in 0.5% BSA in PBS and once with 0.5% BSA in PBS. After washing, the cells were measured in BD Fortessa flow cytometer. Data of both panels was further analyzed using FACSDiva during acquisition and Flowjo for post-analysis.

**Table 1:** Antibodies used for flow cytometry analysis.

| Antibody | Manufacturer | Catalog number | Dilution |
| --- | --- | --- | --- |
| **LDnearIR-FVD780** | Thermo Fisher Scientific | 65-0864-14 | 1:1000 |
| **CD163-BV785** | Biolegend | 333631 | 1:100 |
| **CD8-PE** | Biolegend | 344706 | 1:100 |
| **NGFR-APC** | Biolegend | 345107 | 1:100 |
| **CD146-APC** | Biolegend | 361015 | 1:100 |
| **CD137-PECy7** | Biolegend | 309818 | 1:100 |
| **PD1-BV605** | Biolegend | 329924 | 1:100 |
| **Siglec-1-BV421** | Biolegend | 346018 | 1:100 |
| **CD8-BV421** | BD Biosciences | 562428 | 1:100 |
| **Siglec-1-Fc-AF488** | Invitrogen | A11013 | 1:133 |
| **IFNg-PE** | BD Pharmingen | 554701 | 1:100 |
| **TNF-PerCP** | Biolegend | 502924 | 1:100 |

### Spatial transcriptomic analysis

We performed spatial transcriptomic analysis as described earlier (Coccimiglio, Ibáñez-Molero et al., in revisions). Briefly, we analyzed single-cell spatial transcriptomics data from archived FFPE tumour tissues of four patients with stage III melanoma treated with neoadjuvant immune checkpoint blockade at Amsterdam UMC. Two responders (with major pathological response) and two non-responders were selected. Tissue sections were processed using the Xenium platform (10x Genomics), including RNA extraction, probe hybridization, imaging, and signal decoding to generate spatial gene expression data. Data were analyzed in R using Seurat, with quality filtering, integration, clustering, and manual cell-type annotation based on lineage markers. Low-quality and mixed clusters were excluded, and spatial re-clustering was performed where necessary.

### Statistical analysis

The statistical tests used are stated in each figure. GraphPad and R were used for the analysis.

## Acknowledgements

We thank our colleagues in the department of Molecular Biology and Immunology (MCBI) for fruitful discussions and input. In particular, we thank Alsya Affandi and Joke de Haan for their input on SIGLEC1 blocking antibody. We also like to acknowledge the Flow Cytometry Facility at MCBI and Pathology Facility at Amsterdam Medical Center (AMC) for their technical support in the study, in particular Sara Rohani, Juan García-Vallejo and Sara García-García. We thank our financial support from KWF 2024-3 EXPL / 16684, SPINOZA NWO SPI-93-538, European Union Horizon 2020, Marie Skłodowska-Curie Actions Grant agreement No. 956758, GLYTUNES Consortium, Amsterdam Institute for Immunology and Infectious Diseases and Oncode Accelerator, a Dutch National Growth Fund project under grant number NGFOP2201.

## Author contributions

S.I.M., Y.v.K conceived the study and designed the experiments. S.I.M., M.C., B.O.S., M. B. R., performed the experiments. S.I.M. performed bioinformatics analysis. S.W. and C.B. collected and analyzed single-cell RNA sequencing data from ICB-treated melanoma cohort. S.I.M., G.C., M.L., T.d.G., and Y.v.K. designed and analyzed the melanoma cohort for spatial transcriptomics. S.I.M. and Y.v.K. wrote the manuscript. All authors read and approved the manuscript. The project was supervised by Y.v.K.

## Data availability statement

Processed spatial transcriptomics data are available upon request to authors and they will be deposited in a public data repository. All other raw data are available upon request.

## Code availability statement

Code is available upon request to authors and it will be deposited in github.

## EXTENDED DATA FIGURE AND FIGURE LEGENDS

**Extended Data Figure 1.**
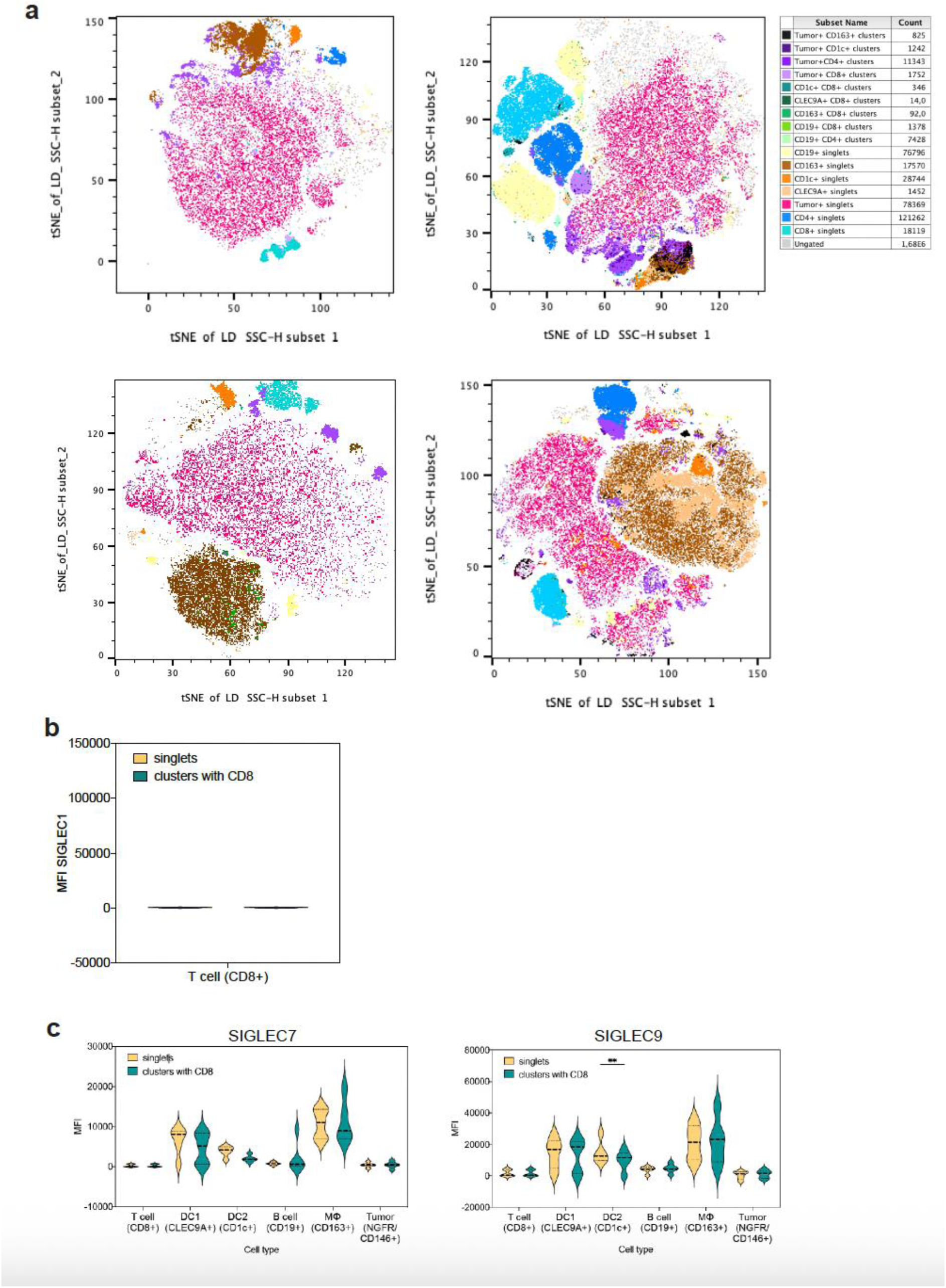
a) tSNE singlets and clusters found in four melanoma patients. b) SIGLEC1 expression in singlets or clustered CD8+ T cells. 2way ANOVA test used for statistical analysis. c) SIGLEC7 and SIGLEC9 expression in different APC subtypes as singlets or clusters with CD8+ T cells. 2way ANOVA test used for statistical analysis.

**Extended Data Figure 2.**
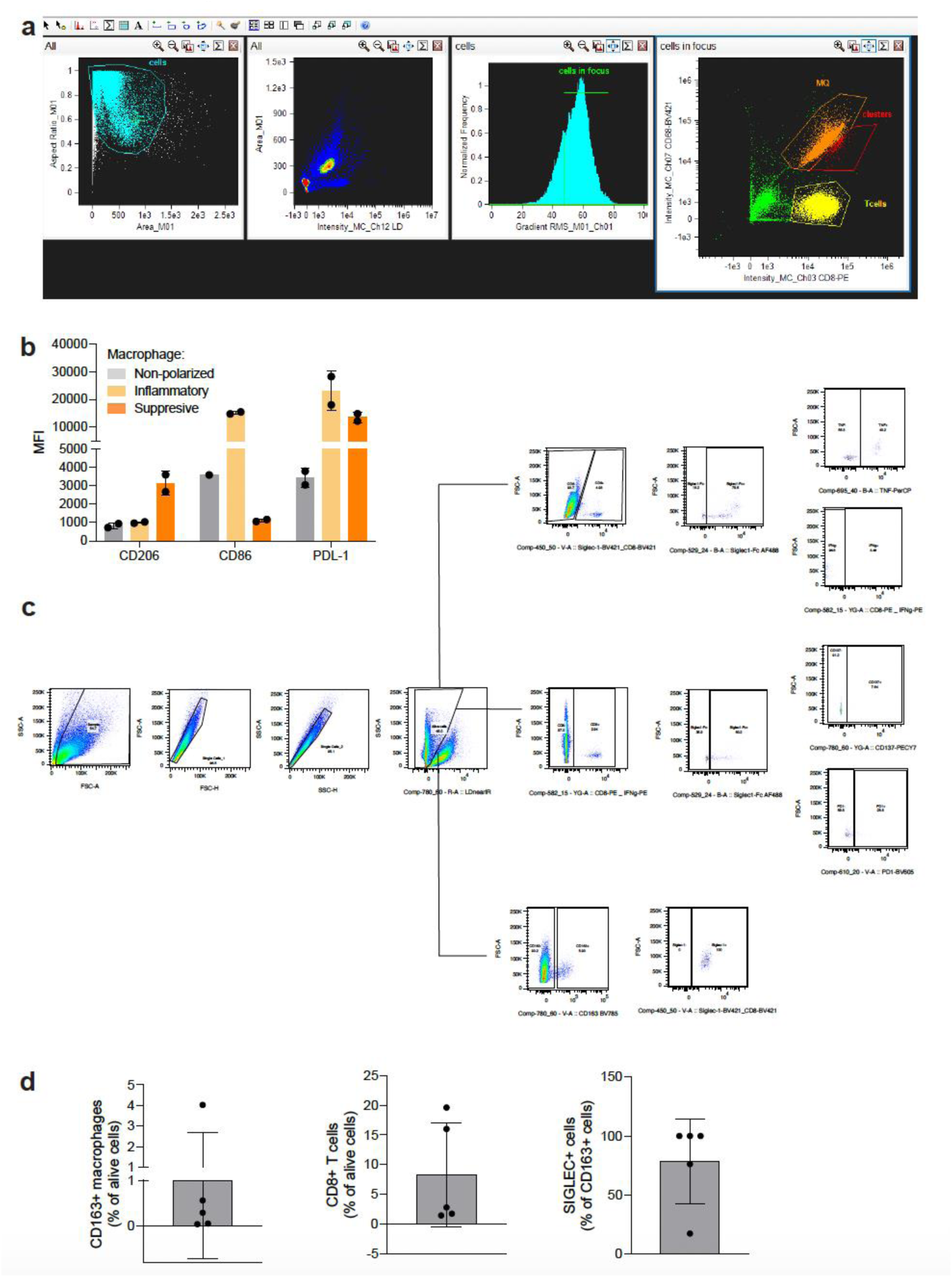
a) Imaging flow cytometry gating strategy of experiment in Fig. 2b-d. b) Median fluorescence intensity of macrophage phenotype markers (CD206, CD86, PDL-1) from experiment in Fig. 2b-f. c) Flow cytometry gating strategy of experiment in Fig. 3b-c. d) Frequencies of CD8^+^ T cells and macrophages from alive cells, and SIGLEC1+ macrophages from macrophage population.

**Extended Data Figure 3.**
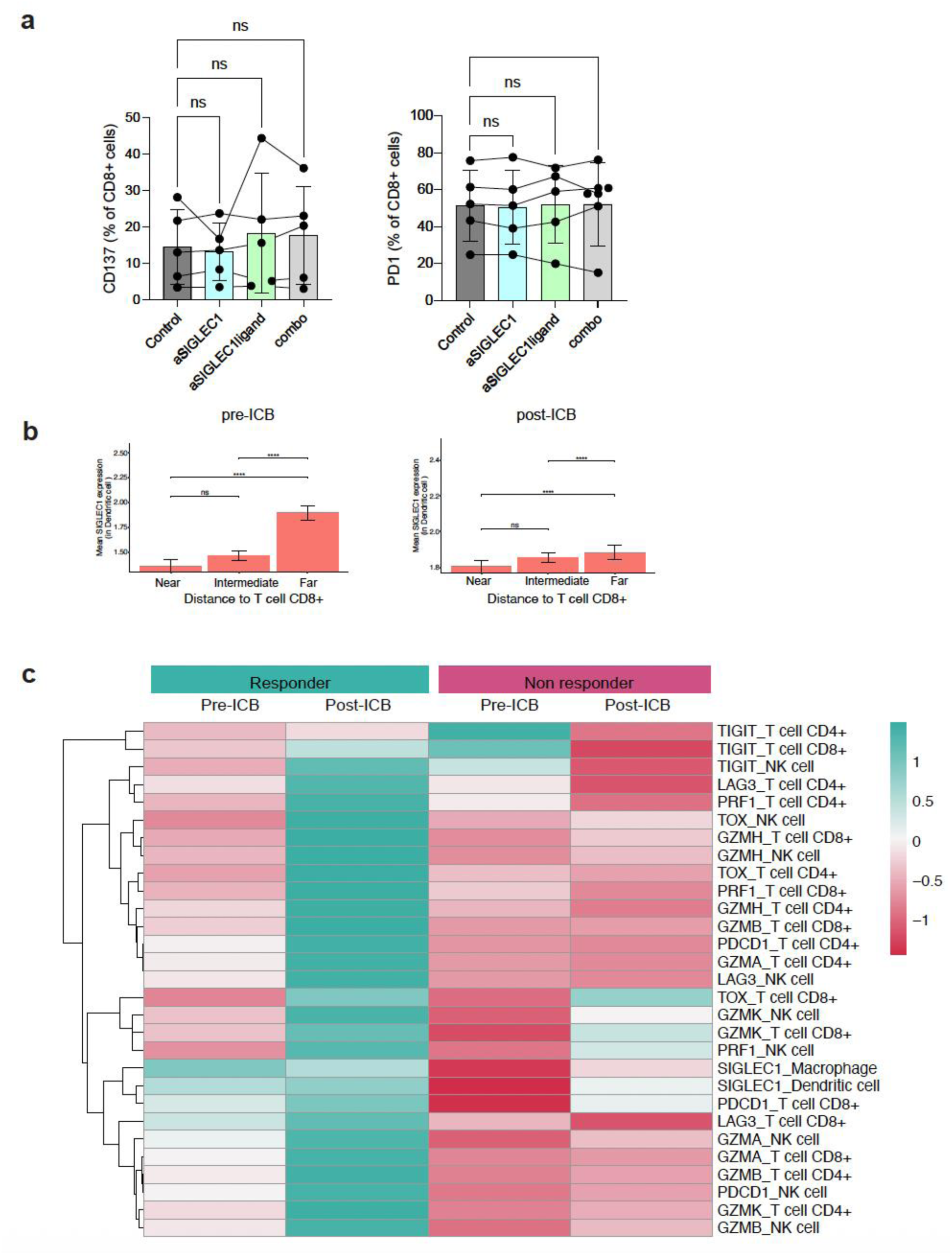
a) Quantification of frequency of PD1 and CD137 positive CD8+ T cells from Fig. 3b. Lognormal RM one-way ANOVA test used for statistical analysis. b) Mean SIGLEC1 expression in DCs relative to distance groups from the closest target cell. “Near” corresponds to the first tercile, “Intermediate” to the second tercile, and “Far” to the third tercile of all distances calculated. Bar plots show the average SIGLEC1 expression per group with standard deviation. Wilcoxon test with Bonferroni correction used for statistical analysis. Analysis performed on pre-ICB samples (left) and post-ICB samples. c) Heatmap showing gene expression of the indicated gene in the indicated cell type (eg. SIGLEC1 gene expression in macrophage cells). Unsupervised row clustered. Blue indicates higher expression, rede indicates lower expression.

## References

1. Blank, C. U. et al. Neoadjuvant Nivolumab and Ipilimumab in Resectable Stage III Melanoma. N. Engl. J. Med. 391, 1696–1708 (2024).

2. Ladányi, A. et al. Density of DC-LAMP+ mature dendritic cells in combination with activated T lymphocytes infiltrating primary cutaneous melanoma is a strong independent prognostic factor. Cancer Immunol. Immunother. 56, 1459–1469 (2007).

3. Yan, C. et al. Spatial distribution of tumor-infiltrating T cells indicated immune response status under chemoradiotherapy plus PD-1 blockade in esophageal cancer. Front. Immunol. 14, 1138054 (2023).

4. Elomaa, H. et al. Spatially resolved multimarker evaluation of CD274 (PD-L1)/PDCD1 (PD-1) immune checkpoint expression and macrophage polarisation in colorectal cancer. Br. J. Cancer 128, 2104–2115 (2023).

5. Homicsko, K. et al. PD-1-expressing macrophages and CD8 T cells are independent predictors of clinical benefit from PD-1 inhibition in advanced mesothelioma. J. Immunother. Cancer 11, e007585 (2023).

6. Wang, J. et al. Multiplexed immunofluorescence identifies high stromal CD68+PD-L1+ macrophages as a predictor of improved survival in triple negative breast cancer. Sci. Rep. 11, 21608 (2021).

7. Kersten, K. et al. Spatiotemporal co-dependency between macrophages and exhausted CD8+ T cells in cancer. Cancer Cell 40, 624–638.e9 (2022).

8. Van Der Leun, A. M., Thommen, D. S. & Schumacher, T. N. CD8+ T cell states in human cancer: insights from single-cell analysis. Nat. Rev. Cancer 20, 218–232 (2020).

9. Komohara, Y., Jinushi, M. & Takeya, M. Clinical significance of macrophage heterogeneity in human malignant tumors. Cancer Sci. 105, 1–8 (2014).

10. Marcovecchio, P. M., Thomas, G. & Salek-Ardakani, S. CXCL9-expressing tumor-associated macrophages: new players in the fight against cancer. J. Immunother. Cancer 9, e002045 (2021).

11. Cheng, S. et al. A pan-cancer single-cell transcriptional atlas of tumor infiltrating myeloid cells. Cell 184, 792–809.e23 (2021).

12. De Logu, F. et al. Spatial Proximity and Relative Distribution of Tumor-Infiltrating Lymphocytes and Macrophages Predict Survival in Melanoma. Lab. Invest. 103, 100259 (2023).

13. Chen, J. H., et al. Spatial Analysis of Human Lung Cancer Reveals Organized Immune Hubs Enriched for Stem-like CD8 T Cells and Associated with Immunotherapy Response. http://biorxiv.org/lookup/doi/10.1101/2023.04.04.535379 (2023) doi:10.1101/2023.04.04.535379.

14. Scott, A. C. et al. TOX is a critical regulator of tumour-specific T cell differentiation. Nature 571, 270–274 (2019).

15. Kubota, K. et al. CD163+CD204+ tumor-associated macrophages contribute to T cell regulation via interleukin-10 and PD-L1 production in oral squamous cell carcinoma. Sci. Rep. 7, 1755 (2017).

16. Boissonnas, A. et al. CD8+ Tumor-Infiltrating T Cells Are Trapped in the Tumor-Dendritic Cell Network. Neoplasia 15, 85-IN26 (2013).

17. Peranzoni, E. et al. Macrophages impede CD8 T cells from reaching tumor cells and limit the efficacy of anti–PD-1 treatment. Proc. Natl. Acad. Sci. 115, (2018).

18. Ibáñez-Molero, S. et al. Tumour-reactive heterotypic CD8 T cell clusters from clinical samples. Nature 649, 467–476 (2026).

19. Prenzler, S. et al. The role of sialic acid-binding immunoglobulin-like-lectin-1 (siglec-1) in immunology and infectious disease. Int. Rev. Immunol. 42, 113–138 (2023).

20. Klaas, M., Dubock, S., Ferguson, D. J. P. & Crocker, P. R. Sialoadhesin (CD169/Siglec-1) is an extended molecule that escapes inhibitory cis-interactions and synergizes with other macrophage receptors to promote phagocytosis. Glycoconj. J. 40, 213–223 (2023).

21. Miller, M. J., Wei, S. H., Parker, I. & Cahalan, M. D. Two-Photon Imaging of Lymphocyte Motility and Antigen Response in Intact Lymph Node. Science 296, 1869–1873 (2002).

22. Monks, C. R. F., Freiberg, B. A., Kupfer, H., Sciaky, N. & Kupfer, A. Three-dimensional segregation of supramolecular activation clusters in T cells. Nature 395, 82–86 (1998).

23. Dustin, M. L., Chakraborty, A. K. & Shaw, A. S. Understanding the Structure and Function of the Immunological Synapse. Cold Spring Harb. Perspect. Biol. 2, a002311–a002311 (2010).

24. Depoil, D. et al. Immunological Synapses Are Versatile Structures Enabling Selective T Cell Polarization. Immunity 22, 185–194 (2005).

25. Dekkers, J. F. et al. Uncovering the mode of action of engineered T cells in patient cancer organoids. Nat. Biotechnol. 41, 60–69 (2023).

26. Affandi, A. J. et al. Selective tumor antigen vaccine delivery to human CD169^+^ antigen-presenting cells using ganglioside-liposomes. Proc. Natl. Acad. Sci. 117, 27528–27539 (2020).

27. Van Dinther, D. et al. Comparison of Protein and Peptide Targeting for the Development of a CD169-Based Vaccination Strategy Against Melanoma. Front. Immunol. 9, 1997 (2018).

28. Prenzler, S. et al. The role of sialic acid-binding immunoglobulin-like-lectin-1 (siglec-1) in immunology and infectious disease. Int. Rev. Immunol. 42, 113–138 (2023).

29. Van Den Berg, T. K., et al. Sialoadhesin on macrophages: its identification as a lymphocyte adhesion molecule. J. Exp. Med. 176, 647–655 (1992).

30. Azevedo, C. M. et al. Reprogramming CD8+ T-cell Branched *N-* Glycosylation Limits Exhaustion, Enhancing Cytotoxicity and Tumor Killing. Cancer Immunol. Res. 13, 1655– 1673 (2025).

31. Büll, C. et al. Sialic Acid Blockade Suppresses Tumor Growth by Enhancing T-cell– Mediated Tumor Immunity. Cancer Res. 78, 3574–3588 (2018).

32. Affandi, A. J. et al. CD169 Defines Activated CD14+ Monocytes With Enhanced CD8+ T Cell Activation Capacity. Front. Immunol. 12, 697840 (2021).

33. Asano, K. et al. CD169-Positive Macrophages Dominate Antitumor Immunity by Crosspresenting Dead Cell-Associated Antigens. Immunity 34, 85–95 (2011).

34. Ohnishi, K. et al. CD 169-positive macrophages in regional lymph nodes are associated with a favorable prognosis in patients with colorectal carcinoma. Cancer Sci. 104, 1237– 1244 (2013).

35. Ohnishi, K. et al. Prognostic significance of CD 169-positive lymph node sinus macrophages in patients with endometrial carcinoma. Cancer Sci. 107, 846–852 (2016).

36. Izquierdo-Useros, N. et al. Siglec-1 Is a Novel Dendritic Cell Receptor That Mediates HIV-1 Trans-Infection Through Recognition of Viral Membrane Gangliosides. PLoS Biol. 10, e1001448 (2012).

37. Baars, A. et al. Skin tests predict survival after autologous tumor cell vaccination in metastatic melanoma: Experience in 81 patients. Ann. Oncol. 11, 965–970 (2000).

